# Activation of the TRIF pathway and downstream targets results in the development of precancerous lesions during infection with *Helicobacter*

**DOI:** 10.1101/2023.06.04.543598

**Authors:** Prerna Bali, Ivonne Lozano-Pope, Jonathan Hernandez, Monica V Estrada, Maripat Corr, Michael A. Turner, Michael Bouvet, Christopher Benner, Marygorret Obonyo

## Abstract

*Helicobacter pylori* (*H. pylori)* infection is an established cause of many digestive diseases, including gastritis, peptic ulcers, and gastric cancer. However, the mechanism by which infection with *H. pylori* causes these disorders is still not clearly understood. This is due to insufficient knowledge of pathways that promote *H. pylori*-induced disease progression. We have established a *Helicobacter*-induced accelerated disease progression mouse model, which involves infecting mice deficient in the myeloid differentiation primary response 88 gene (*Myd88^-/-^*) with *H. felis*. Using this model, we report here that that progression of *H. felis*-induced inflammation to high-grade dysplasia was associated with activation of type I interferon (IFN-I) signaling pathway and upregulation of related downstream target genes, IFN-stimulated genes (ISGs). These observations were further corroborated by the enrichment of ISRE motifs in the promoters of upregulated genes. Further we showed that *H. felis*-induced inflammation in mice deficient in Toll/interleukin-1 receptor (TIR)-domain-containing adaptor inducing interferon-β (TRIF, *Trif^Lps^*^2^) did not progress to severe gastric pathology, indicating a role of the TRIF signaling pathway in disease pathogenesis and progression. Indeed, survival analysis in gastric biopsy samples from gastric cancer patients illustrated that high expression of *Trif* was significantly associated with poor survival in gastric cancer.

## Introduction

*Helicobacter pylori* (*H. pylori)* infection is the main risk factor for development of gastric cancer and this bacterium is classified as a group I carcinogen^1,2^. Worldwide, gastric cancer remains a significant global health burden as one of the leading causes of cancer-related deaths and one of the most recalcitrant malignancies^3,4^. With half of the world’s population infected with *H. pylori*, this bacterium contributes significantly to the worldwide disease burden.^2,5^ Infection with this organism is known to cause chronic inflammation, which eventually progresses to preneoplasia lesions (atrophy, intestinal metaplasia, dysplasia), and finally to gastric cancer, a process commonly known as the Correa’s cascade.^6^ We previously reported that infection of mice deficient in myeloid differentiation primary response gene 88 (MyD88, *Myd88*^-/-^) with *H. felis* resulted in rapid development of gastric pathology that progressed to high-grade gastric dysplasia^7^, which is considered gastric cancer *in situ*^8,9^. However, the mechanisms leading to this phenotype remained undetermined although the data indicated that the loss of MyD88 triggered a hyperactivation of the alternate Toll/interleukin-1 receptor (TIR) domain-containing adaptor inducing interferon-β (TRIF)-dependent pathway leading the accelerated gastric cancer progression in response to *H. felis* infection^7,10^. A recent study citing this work reported that the TLR4-TRIF-type I interferon (IFN-I) via production of IFN-γ was important for *H. sius*-induced disease development in a mucosa-associated lymphoid tissue (MALT) lymphoma mouse model ^11^. MyD88 and TRIF are the main adaptor proteins that mediate toll-like receptor (TLRs) signals^12^. TRIF signaling diverges into two downstream pathways, one that shares downstream effectors with MyD88 and the other that is MyD88 independent, TRIF/IRF3 (IFN regulatory factor 3)^13–15^, the latter which leads to downstream induction of type I IFN genes and IFN-inducible genes.

A role of the TRIF-mediated arm of TLR signaling in *Helicobacter*-induced disease remains less studied except for our prior suggested observation^7,10^ and a recent study that used a *Helicobacter*-induced MALT lymphoma model^11^. In addition, it remains unclear when gastric epithelial changes occur following *Helicobacter* infection. Furthermore, the role of related downstream target genes on gastric cancer tumorigenesis, comprised primarily of IFN-stimulated genes (ISGs), also remains unknown.

In this study, we used our accelerated animal model for gastric cancer and patient clinical gastric specimens to show that activation of the TRIF-IFN-I signaling pathway promotes *Helicobacter*-induced disease progression to severe gastric pathology that ultimately leads to the development of gastric cancer. This pathway could therefore be targeted to intercept the neoplastic process with a goal of preventing the development of gastric cancer.

## Material and methods

### Animals

Six- to ten-week-old wild type (WT) (n=42), MyD88 deficient (*Myd88^-/-^*) (n=47), TRIF deficient (*Trif*^Lps2^) (n=46), double knockout (*Myd88^-/-^* and *Trif*^Lps2^, DKO) (n=37) mice in the C57BL/6 background were used in this study. WT mice were purchased from The Jackson Laboratory (Bar Harbor, ME). *Myd88*^-/-^ mice were from our breeding colony originally provided by Dr. Akira (Osaka University, Japan) and backcrossed 10 times onto a C57BL/6 background, bred, and maintained at our facility. The DKO mice were bred in our facility by crossing *Myd88^-/-^* and *Trif*^Lps2^. All mice were housed together before infection with *H. felis* and for the duration of the study for each genotype. The institutional Animal Care and Use Committee at the University of California, San Diego, approved all animal procedures and performed using accepted veterinary standards.

### Bacterial growth conditions

*Helicobacter felis*, strain CS1 (ATCC 49179) was purchased from American Type Culture collection (Manassas, VA). *H. felis* was routinely maintained on solid medium, Columbia agar (Becton Dickinson, MD) supplemented with 5% laked blood under microaerophilic conditions (5% O2, 10% CO2, 85% N2) at 37 °C and passaged every 2–3 days as described previously ^7,16–18^. Prior to mouse infections, *H. felis* was cultured in liquid medium, brain heart infusion broth (BHI, Becton Dickinson) supplemented with 10% fetal calf serum and incubated at 37 °C under microaerophilic conditions for 48 h. Spiral bacteria were enumerated using a Petroff-Hausser chamber before infections.

### Mouse infections

This study used a well-characterized cancer mouse model, which involves infecting C57BL/6 mice with *H. felis* (strain CS1), a close relative of the human gastric pathogen *H. pylori*. Mice were inoculated with 10^9^-organisms in 300 μL of BHI by oral gavage three times at 2-day intervals as previously described ^7,16–18^. Control mice received BHI only. At 1 month, 3 months and 6 months post infection, mice were euthanized, and the stomachs removed under aseptic conditions. The stomach was cut longitudinally, and tissue sections were processed for RNA extraction and histopathology. All *H. felis*-infected mice were confirmed colonized by assessing expression of the flagella filament B (*flab*) gene by real-time PCR as described in our previous studies ^7^. There was no significant difference in expression of the *flabB* gene between all infected mice groups.

### Histology

Mouse stomachs were processed for histology as described in our previous studies^7,19^. Briefly, longitudinal sections of stomach tissue from each mouse were fixed in neutral buffered 10% formalin and embedded in paraffin, and 5 µm sections were stained with hematoxylin and eosin (H&E). Gastric histopathologies, including inflammation, epithelial defects, gland atrophy, hyperplasia, intestinal metaplasia, and dysplasia, were scored double-blind by a pathologist (Dr. Estrada) for histologic disease severity on an ascending scale from 0 (no lesions) to 4 (severe lesions) as described in our previous studies ^7,19^ using criteria outlined by Rogers et al.^20^. Specifically, the definition of the histological characteristics of the parameters we evaluated were as follows: Inflammation; an infiltration of inflammatory cells into the mucosa and submucosa, oxyntic gland atrophy: loss of oxyntic glands (which contain parietal cells and chief cells) in the gastric corpus, hyperplasia; elongation of gastric glands due to increased numbers of surface cells (foveolar cells), intestinal metaplasia; replacement of gastric epithelium by intestinal type gastric epithelium, and dysplasia; abnormal cellular and glandular maturation of the epithelium, with score of 3 considered carcinoma *in situ*. ^20^

### Patient gastric biopsy samples

De-identified snap frozen twelve human gastric biopsy samples were obtained from the UCSD’s Biorepository. All patients provided written informed consent and were followed up. Details about the patients and their gastric tumor characteristics are provided in Table S1^21^. RNA was extracted from each biopsy tissue and processed for real-time PCR as described below.

### RNA extraction from mouse and human gastric tissue samples

RNA was extracted from gastric tissue obtained from *H. felis*-infected and uninfected WT, *Myd88*^-/-^, *Trif^Lps^,* and DKO mice. Stomach tissue sections were analyzed at different time points of 1 month, 3 months, and 6 months. Total RNA from each mouse gastric tissue section was homogenized in 1 ml of TRIzol reagent (Invitrogen, Carlsbad, CA) according to the manufacturer’s instructions and RNA was extracted using Direct-zol RNA mini kit (Zymo Research, Irvine CA) according to the manufacturer’s instructions and stored in - 70°C till further use. The quality of RNA was determined using a Nanodrop system (Thermo Fisher, Waltham MA) by reading absorbance levels at 260 and 280 nm. RNA was utilized downstream for RNASeq analysis and qPCR. Total RNA was also extracted from human biopsy tissue samples using this method.

### cDNA synthesis and quantitative real-time RT-PCR

Real-time RT-PCR (reverse transcription polymerase chain reaction) was performed as previously described in our studies^7,10,18,19^. We determined the expression of select genes in the TRIF-IFN-I pathway including, indoleamine 2,3-dioxygenase 1 (*Ido1*), interferon gamma-induced GTPase (*Igtp*), guanylate binding protein 2 (*Gbp2*), transcription factor Interferon regulatory factor 1 (*Irf1*), 2’-5’-oligoadenylate synthase 2 (*Oas2*), and chemokine (C-X-C motif) ligand 11 (*Cxcl11*), *Cxcl9*, MX dynamin-like GTPase 1(*Mx1*), proteasome subunit beta type 8 (*psmb8*), immunity-regulated GTPase family M protein 1(*Irgm1*), interferon-induced protein with tetratricopeptide repeats 1 (*Ifit1*), and interferon-stimulated gene 15 (*Isg15*). Samples from mouse stomach tissue sections and human gastric biopsy from gastric cancer patients (UCSD Biorepository) were used in this experiment. RNA isolated from gastric tissue samples (2μg/sample) was reverse transcribed into cDNA using High Capacity cDNA Reverse Transcription Kit (Thermo Fisher, Waltham MA). 1μl of cDNA was used per well in a total of 10μl reaction mix for amplification using the StepOne Real Time PCR (Applied Biosystems, Carlsbad CA). The amplification conditions consisted of an initial cycle of 95°C for 5 minutes followed by 40 cycles of amplification with denaturation as follows: 95°C for 15 sec, 60°C for 20 sec, 72°C for 40 sec. Gene expression levels were normalized to Glyceraldehyde 3-phosphate dehydrogenase (*Gapdh*). The data collected were analyzed using a comparative cycle threshold calculations (ΔΔC_T_, Applied Biosystems) and plotted using GraphPad Prism software. The primers used in this study are listed in Table S2.

### RNA sequencing

RNA concentration and integrity was assessed using an Agilent 2100 Bioanalyzer and each sample assigned RNA integrity number (RIN). Selected samples were processed into RNA-seq libraries using the Illumina TruSeq Stranded Total RNA Sample Prep Kit with Ribo-Zero (Human/Mouse/Rat), which performs ribosomal RNA depletion, fragmentation, cDNA synthesis, and cDNA library preparation. Sequencing was carried out on an Illumina/Solexa HiSeq 2500 sequencer to generate 75-bp single-end reads for all samples at a depth of approximately 40 million reads per sample.

### RNA-seq analysis

RNA-seq reads were aligned were aligned to the mouse genome (GRCm38/mm10) using STAR with default parameters ^22^. Only reads aligning uniquely to a single location in the genome were used for downstream analysis (MAPQ > 10). Gene expression was determined by counting the number of reads overlapping exons for all genes using HOMER’s analyzeRepeats.pl tool and the transcriptome annotation from GENCODE (version M25) ^23^. DESeq2 was used to identify differentially expressed genes from unnormalized read counts^24^. Volcano plot visualization was performed using the EnhancedVolcano package from R/Bioconductor. Gene expression values were normalized for heatmap visualization using DESeq2’s rlog function. Normalized genome browser read density plots were generated by first creating bedGraph files with HOMER and then visualizing them using the UCSC Genome Browser.

### Pathway and motif enrichment analysis

Pathway enrichment analysis was performed with Metascape ^25^, web-based portal using multiple knowledgebases, including the Gene Ontology, to identify overrepresented pathways in sets of regulated genes. Differentially expressed genes with a fold change of at least 2 and an adjusted p-value of 0.05 were used for Metascape analysis. The reported Metascape pathway enrichment results are corrected for multiple hypothesis testing. Transcription factor motif analysis of mouse stomach tissue sections (RNA-seq) was performed using HOMER, a software suite for DNA motif and Next-generation sequence (NGS) analysis ^23^. Known transcription factor motifs from the HOMER database were scanned in mouse promoter sequences defined from -300 to +50 bp relative to GENCODE defined transcription start site (TSS) locations. The reported log p-values were corrected for multiple hypothesis testing using the Benjamini–Hochberg procedure.

### Human gastric ISG gene expression

Gene expression levels in gastric tissues were reanalyzed using data Gene Expression Omnibus (GEO) database accession number GSE27411^26^. This data contained microarray gene expression of gastric tissue from individuals with or without *H. pylori* infection^26^. RNA-seq expression values were quantile normalized and visualized as a heatmap using Java Tree View^27^.

### Survival analyses of *Myd88* and *Trif* using the Kaplan-Meier plotter

The Kaplan-Meier plotter bioinformatics platform (https://kmplot.com/analysis/) was used to analyze the effect of *Myd88* and *Trif* expression on overall survival (OS), first progression (FP), and post-progression survival (PPS) in gastric cancer patients using data from the GEO database^28^. This online database contains microarray gene expression data of 1065 gastric cancer patients.

### Statistics

Statistical analysis was performed using GraphPad Prism (La Jolla, CA, USA). Normal distributed data was analyzed using T-test, while non-normally distribute data were analyzed using Mann Whitney U Test. For multiple comparisons, ANOVA with Bonferroni’s correction (normal distribution) or Kruskal-Wallis with Dunn’s comparison test (non-normal distribution) were used. Survival curves and analysis were performed using the Kaplan-Meier method and the hazard ratio (HR) with 95% confidence interval. The patients were divided into high and low expression groups using the best cut-off and statistical differences were tested by the log-rank test. *P*-values ˂0.05 were considered significant. Differentially expressed genes were defined as genes with a fold change of at least 2 and an adjusted p-value of 0.05 as determined by DESeq2.

## Results

### Preneoplastic gastric lesions develop early in the absence of Myd88 signaling in response to *Helicobacter* infection in mice

Our previous in vivo data using *Myd88*^-/-^ mice showed that infection for 25 or 47 weeks with *H. felis* resulted in rapid disease progression with severe gastric pathology ^7^. However, it remained unknown when gastric neoplastic changes were initiated following infection with *H. felis*. Therefore, to determine when these gastric epithelial changes first appear, we profiled the gastric epithelium at earlier time points post infection with *H. felis*. WT and *Myd88*^-/-^ mice in the C57BL/6 background were infected with *H. felis* for 1, 3, or 6 months. Because findings from our previous study^7^ indicated a role for the alternate TRIF pathway in promoting *H. felis*-induced disease progression, we also infected *Trif*^Lps2^ and DKO mice with *H. felis* for 1, 3, and 6 months. Gastric histopathological changes were minimal in *Trif^Lps^*^2^ mice in response to infection with *H. felis* compared to WT, *Myd88*^-/-^ and DKO mice (**Fig. 1A**). In contrast, gastric precancerous lesions developed and progressed more rapidly with significantly increased severity in *H. felis*-infected *Myd88*^-/-^ mice compared to *H. felis*-infected WT, *Trif^Lps^*^2^ and DKO mice. To define gastric epithelial changes more closely during *H. felis* infection, histopathology parameters including inflammation, oxyntic gland atrophy, hyperplasia, intestinal metaplasia, and dysplasia were scored on H&E-stained stomach tissue samples by a pathologist blinded to the study. Differences in histopathologic scores of these parameters were evident in *Myd88*^-/-^ mice at 1 month following infection with *H. felis* compared to WT, *Trif^Lps^*^2^, and DKO mice with disease severity increasing with time (**Fig. 1B-1G**). There were no significant gastric histopathologic changes in all uninfected mice (**Fig. S1**.). The earliest significant gastric epithelial changes in these mice occurred at 3 months. Consequently, we performed comparative analyses in mice at 3 months following infection with *H. felis*. Information gathered at this early phase of infection is ideal for identifying early targets as the disease progresses, which is urgently needed given a lack of characteristic symptoms of early disease and as such the disease is usually diagnosed at a terminal stage with bad outcomes for patients.

**Figure 1.**
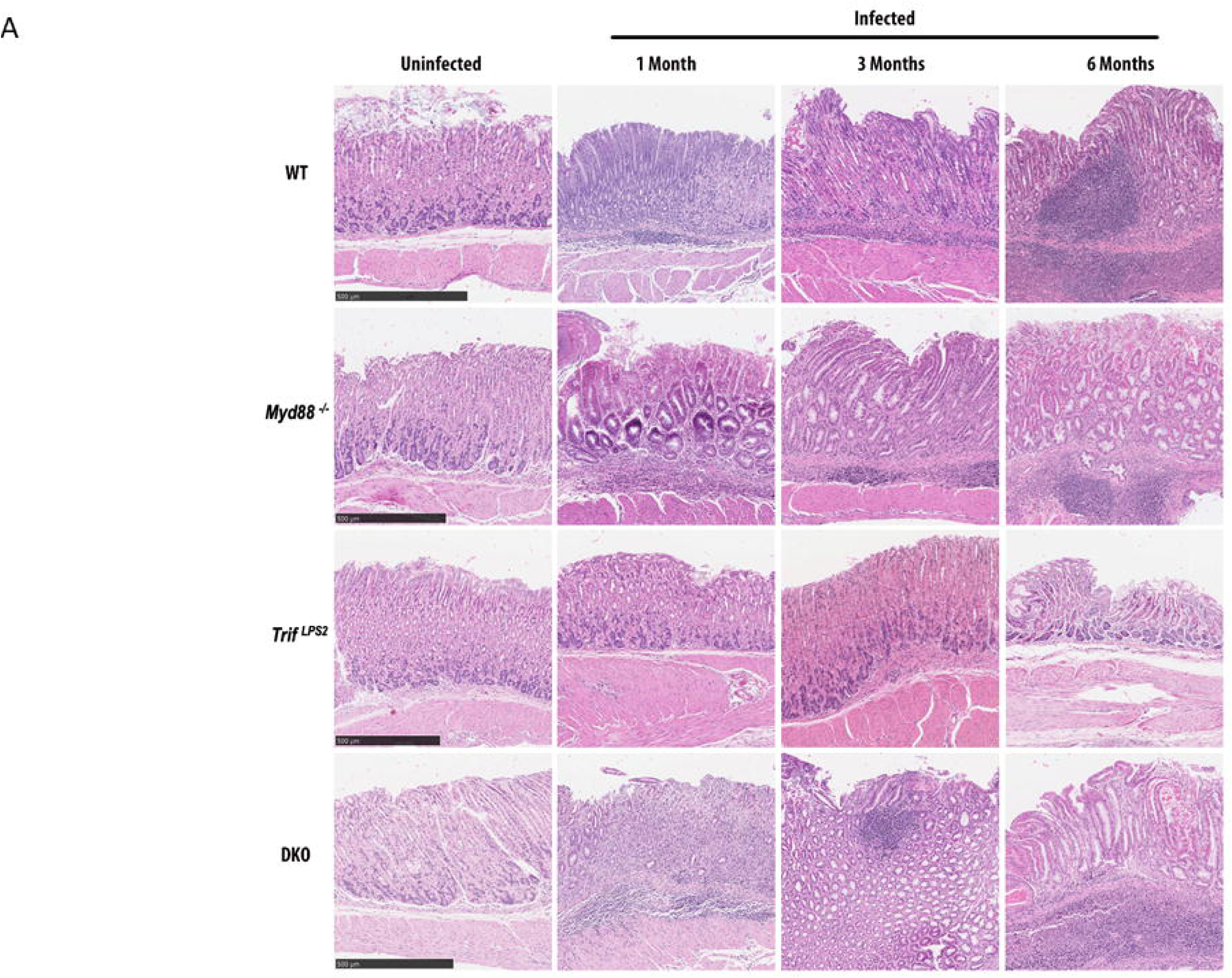
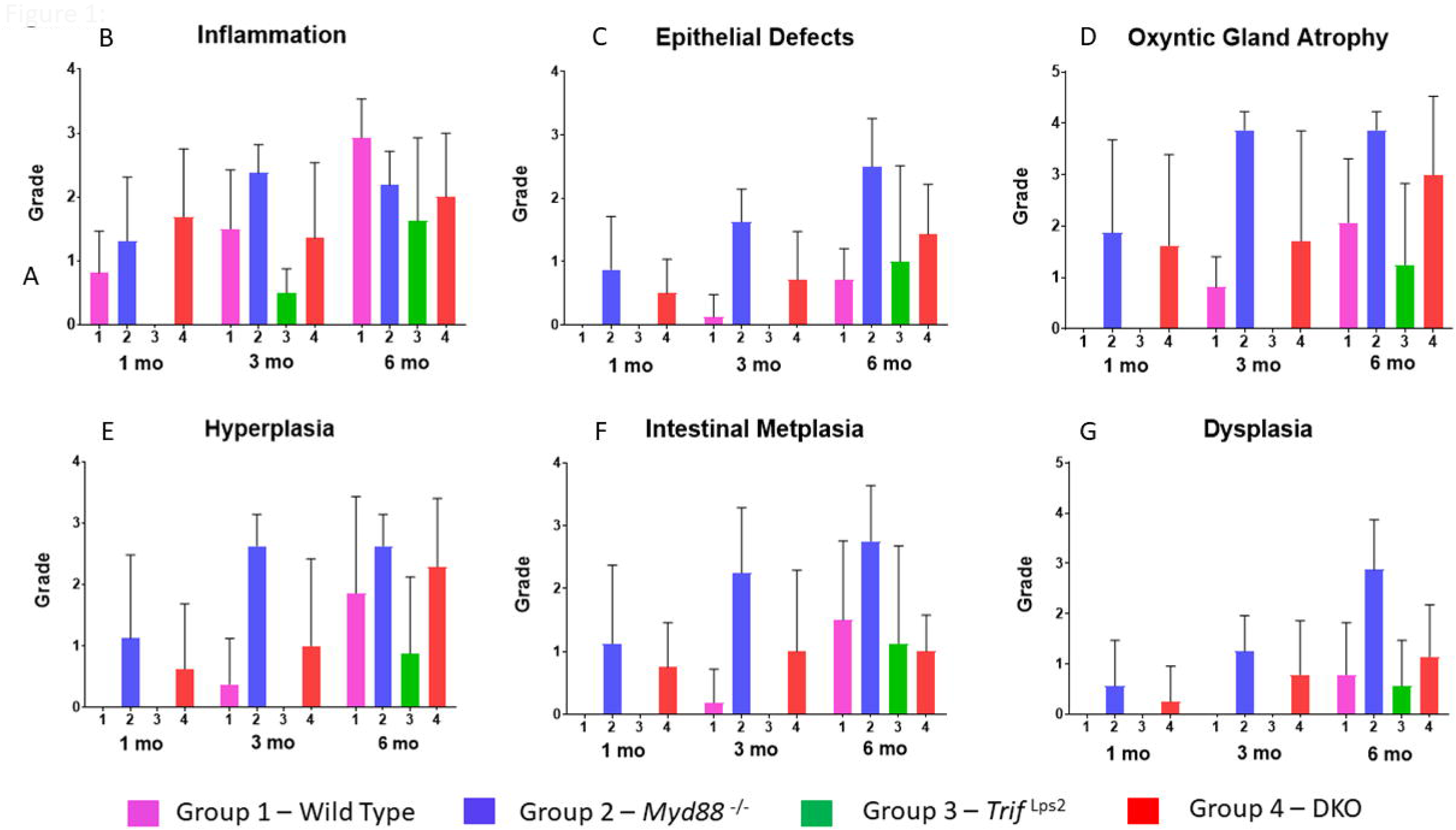
Representative images of hematoxylin & eosin-stained mouse stomach tissue sections from WT, *Myd88*^-/-^, *Trif^Lps2^*, and DKO mice infected with *H. felis* for 1, 3, or, 6 months or left uninfected (A). Mouse stomachs tissue sections were scored for histologic disease severity on an ascending scale from 0 (no lesions) to 4 (severe lesions) for inflammation (B), epithelial defects (C), Oxyntic gland atrophy (D), Hyperplasia (E), Intestinal metaplasia (F), Dysplasia (G).

### *Helicobacter* induced high expression of ISGs in the absence of My88 signaling in mice

Histopathology data indicated significant changes in the gastric epithelium of *Myd88*^-/-^ mice including preneoplastic lesions following infection with *H. felis* as early as 3 months compared to WT, *Trif^Lps^*^2^, or DKO mice. To identify possible molecular mechanisms driving this disease progression in the absence of MyD88 signaling, we compared gene expression in mouse stomachs between WT and *Myd88*^-/-^ mice following infection with *H. felis* for three months using RNA-seq. Analysis of significant differentially expressed genes between *Myd88*^-/-^ and WT mice illustrated that many ISGs were highly expressed in gastric tissue samples from *Myd88*^-/-^ mice (log2 fold change > 1, adj. p-value < 0.05, n=221, **Fig. 2A**), which exhibited severe gastric pathology. Pathway enrichment analysis revealed that these ISGs were the dominant upregulated genes (**Fig. 2B**) and their expression was closely correlated with severe gastric pathology suggesting that their expression plays a role in disease pathogenesis. Some of the ISGs that were upregulated in *Myd88*^-/-^ mice in response to infection with *H. felis* included *Ido1*, *Igtp*, *Gbp2*, *Irf1*, *Cxcl11*, *Cxcl9*, *Cxcl8*, *Mx1*, *Psmb8*, *Irgm1,Zbp1, B2m* (**Fig. 2C**). These ISGs were specifically induced in the gastric tissue from *H. felis*-infected *Myd88*^-/-^ mice and were not observed in gastric tissue samples from *H. felis*-infected WT or uninfected *Myd88*^-/-^ mice.

**Figure 2A.**
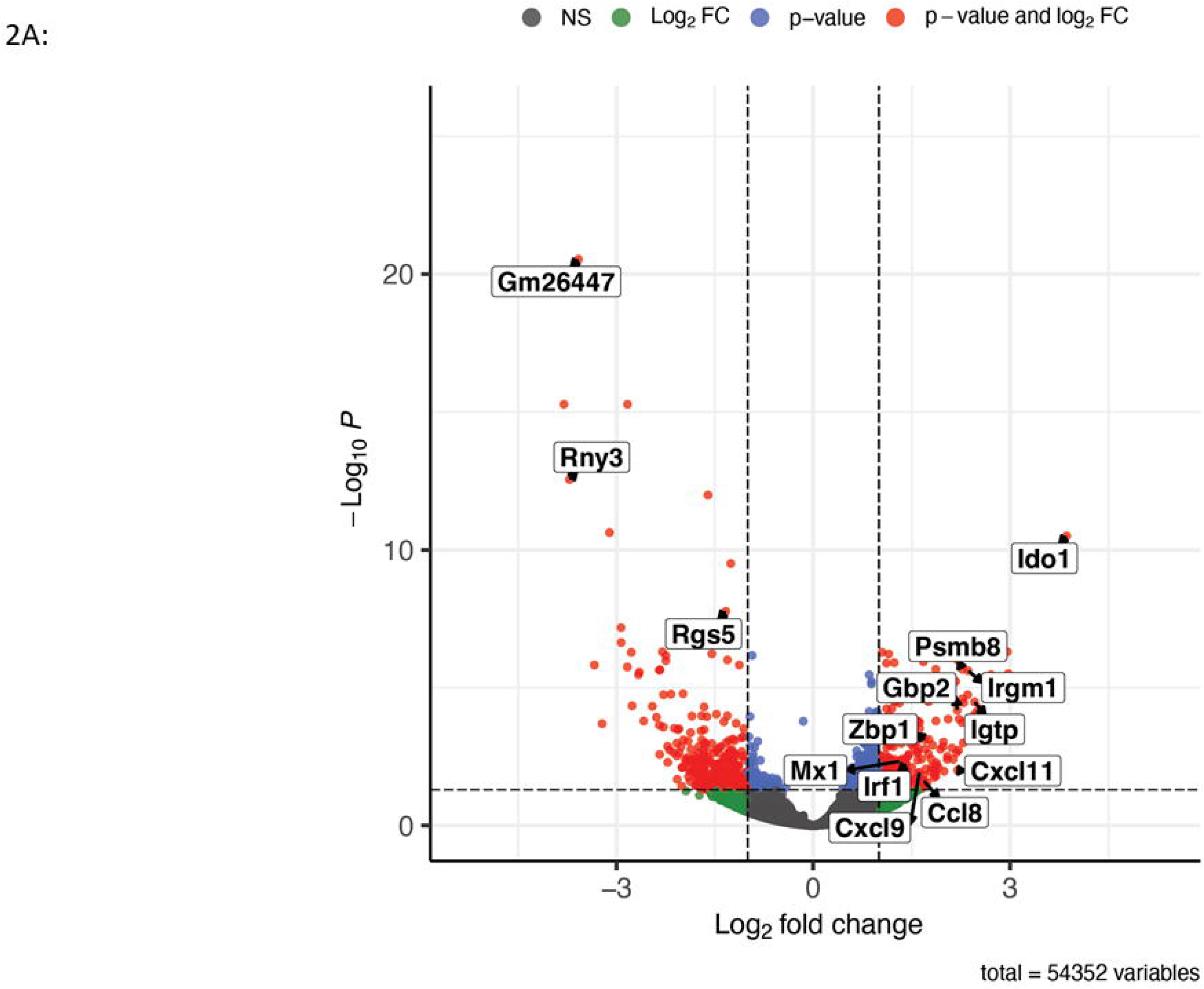
Volcano plot showing genes expressed in mouse gastric tissue. The plot depicts the log2 fold change and adjusted log10 P values for each gene when comparing gene expression values between *felis*-infected WT and *H. felis*-infected *Myd88*^-/-^ mice gastric tissue.

**Figure 2B.**
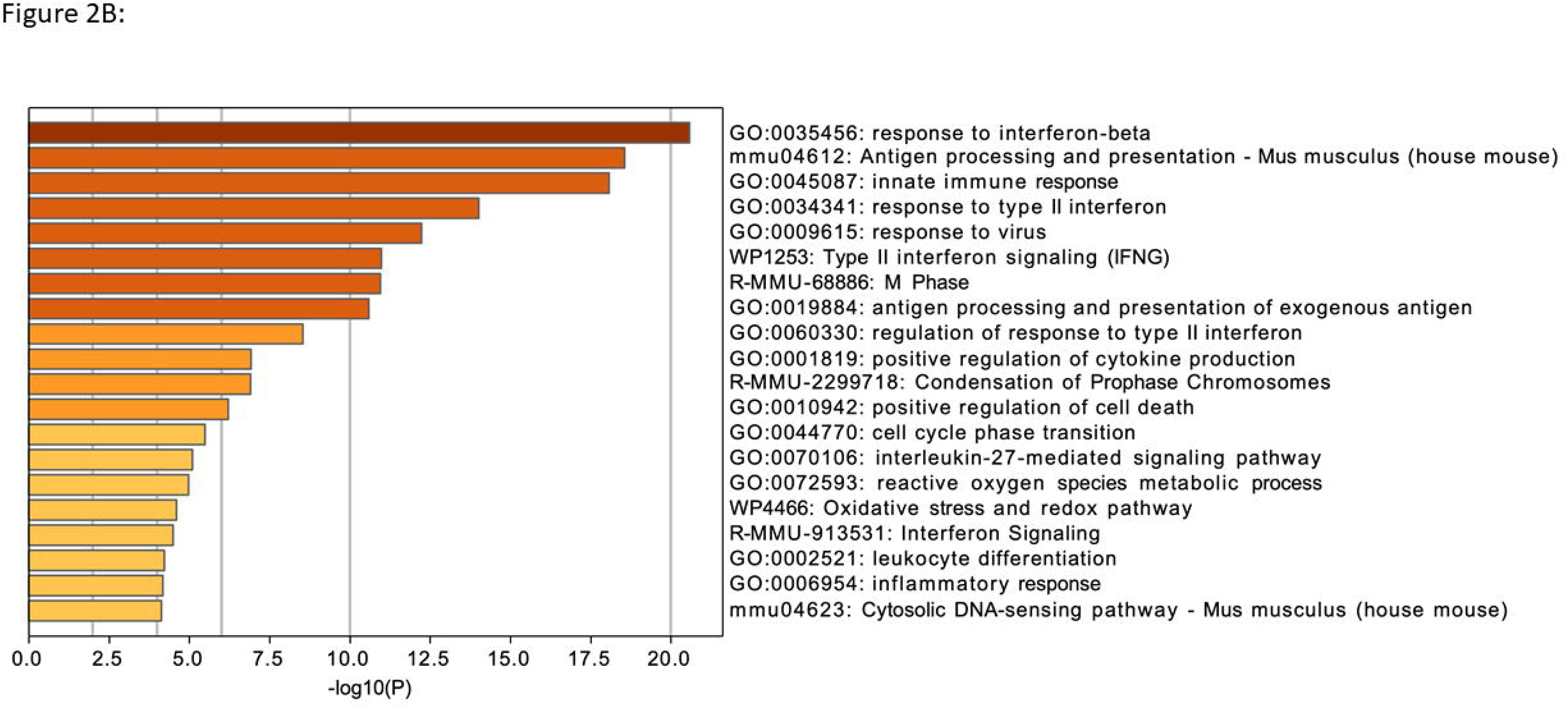
Metascape pathway enrichment analysis showing the top 20 pathways and their adjusted log10 p-value enrichment values in genes that were up-regulated in *H. felis*-infected *Myd88*^-/-^ mice gastric tissue samples relative to *H. felis*-infected WT gastric tissue samples.

**Figure 2C.**
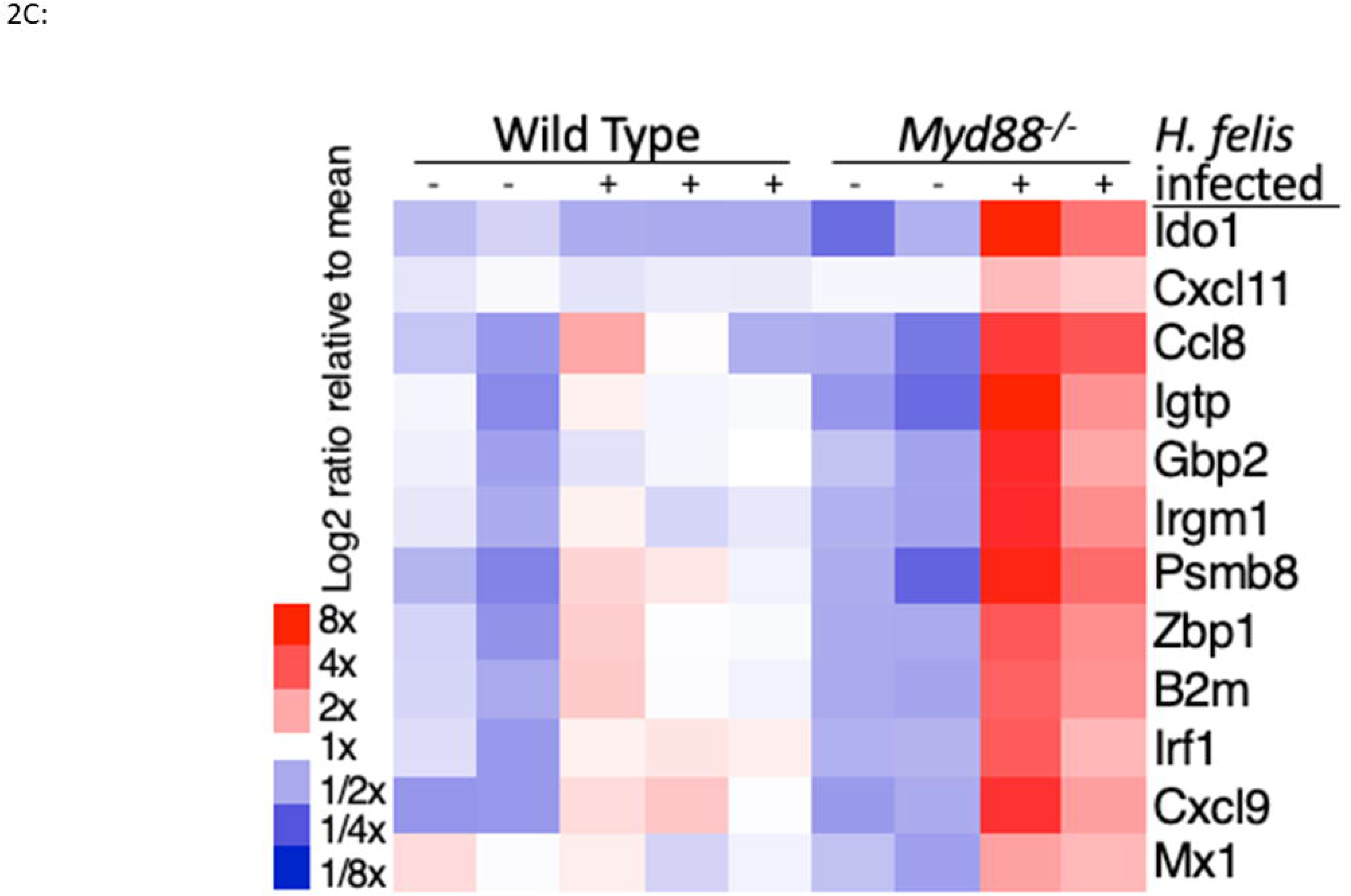
Heat map showing the relative gene expression values for several ISGs across all replicate experiments for uninfected and *H-felis*-infected gastric tissue samples from WT and *Myd88*^-/-^ mice.

**Figure 2D.**
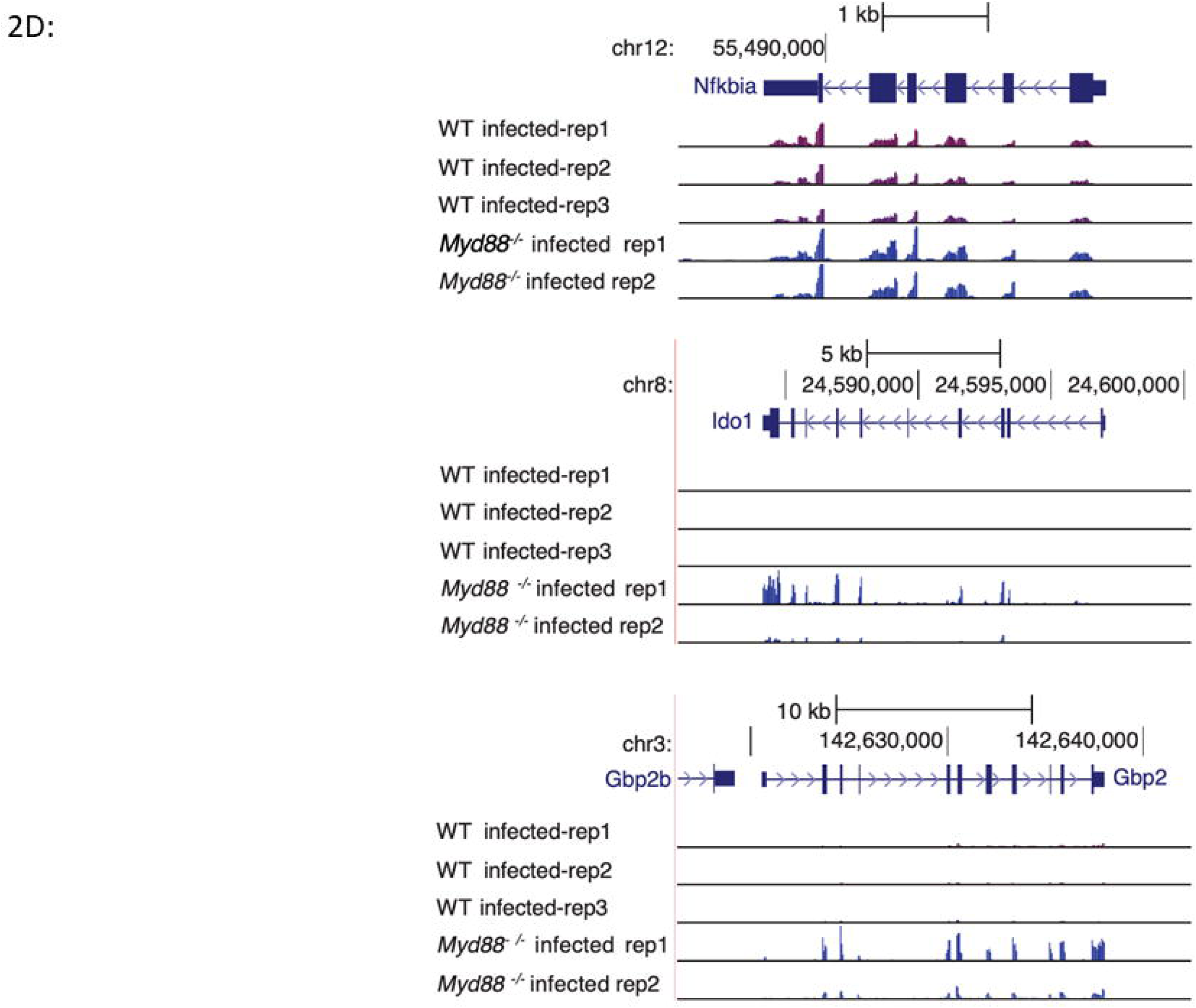
Genome browser tracks displaying the normalized read density for each RNA-seq experiment at the canonical NF-kB target *Nfkbia* locus, and ISG loci *Ido1* and *Gbp2* in gastric tissue samples from WT and *Myd88*^-/-^ mice.

**Figure 2E.**
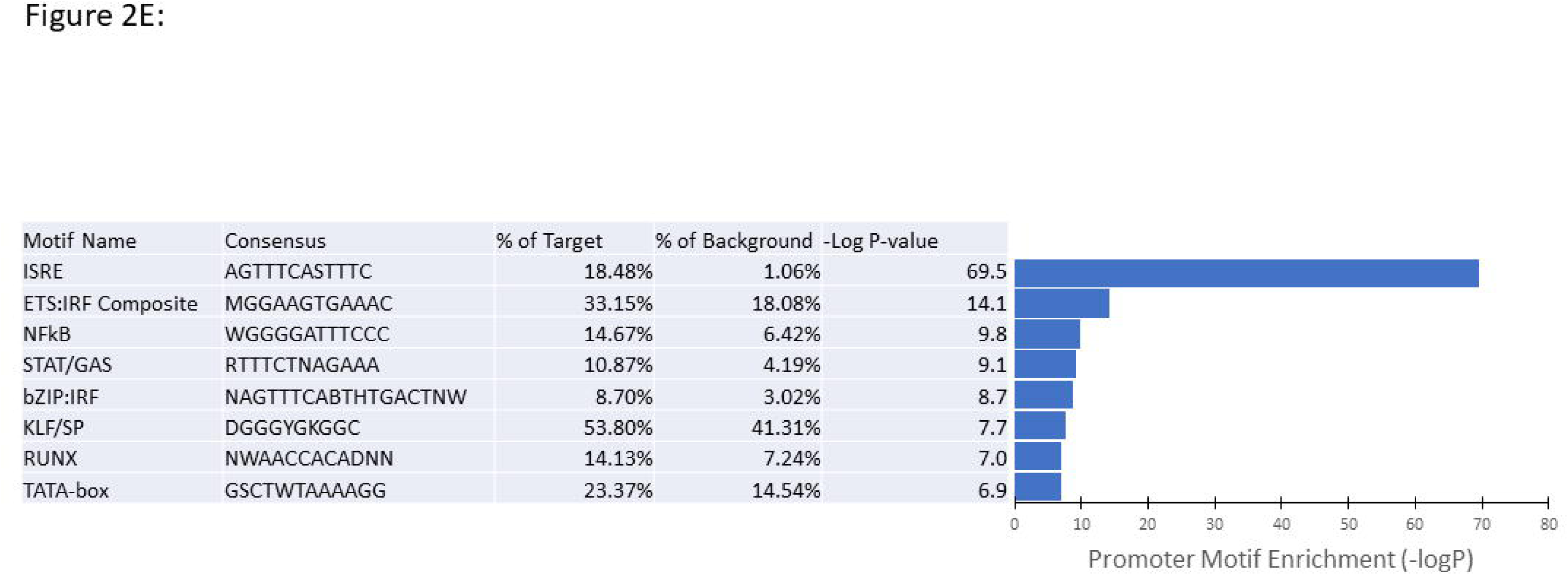
Top 8 non-redundant transcription factor motifs enriched in the promoters of genes up-regulated in gastric tissue from *H. felis*-infected *Myd88*^-/-^ mice relative to *H. felis*-infected WT gastric tissue samples.

We found that upregulation of these ISGs in response to *H. felis* infection was very specific. ISGs are induced in response to activation of the IFN-I pathway.^29^ Thus, comparing upregulation of IFN-I targets with those of transcription nuclear factor-kappa B (NF-κB) in response to *H. felis* infection, showed that while there were minimal differences in upregulation of NF-κB targets between WT and *Myd88*^-/-^ mice, exemplified by the NF-kB target *Nfkbia*, upregulation of IFN-I targets (i.e. ISGs) was only observed in the absence of MyD88 signaling (*Myd88*^-/-^ mice) (**Fig. 2D**). This association of MyD88-deficiency with a heightened ISG profile suggests an activation the TRIF-IFN-I signaling pathway. This specific and strong correlation of ISG upregulation with *H. felis*-induced severe gastric pathology led us to perform promoter motif enrichment analysis of genes induced in the stomach of both WT and *Myd88*^-/-^ mice following infection with *H. felis* to elucidate the possible pathogenic mechanism of these ISGs in *Helicobacter*-induced disease. In line with the upregulation of ISGs in our *H. felis*-infected *Myd88*^-/-^ model, this analysis revealed that of the top 10 enriched transcription factor motifs, the IFN stimulated-response element (ISRE) motif was the most enriched motif in promoters of genes upregulated in *H. felis*-infected *Myd88*^-/-^ infected mice (**Fig. 2E**). This provides evidence that IFN-I regulatory pathways and transcription factors play a role in the severe disease pathology observed in these mice in response to *H. felis* infection. Given that ISGs principally mediate IFN effects^29^, these data therefore, suggest a role of the TRIF-IFN-I pathway in driving severe gastric pathology in response to *H. felis* infection.

### *Helicobacter* induced less disease in the absence of TRIF signaling in mice

Histopathology scoring of gastric tissue from WT, *Myd88*^-/-^, *Trif^Lps^*^2^, and DKO mice following infection with *H. felis* for 3 months revealed that inflammation was induced in all mouse groups but was significantly less severe in *Trif^Lps^*^2^ mice compared to WT, *Myd88*^-/-^, and DKO mice (**Fig. 3A**). While the disease progressed in WT, *Myd88*^-/-^, and DKO mice revealing development of oxyntic gland atrophy following infection with *H. felis*, *Trif^Lps^*^2^ mice did not show evidence of this pathology (**Fig. 3B**). Oxyntic atrophy, which is loss of parietal cells represents a pivotal point in the process of neoplastic lesion development including intestinal metaplasia and dysplasia and ultimately gastric cancer ^30,31^. Indeed, these data show that in the absence of TRIF signaling *Helicobacter*-induced inflammation did not progress to precancerous lesions, including intestinal metaplasia (**Fig. 3C**) and dysplasia **(Fig. 3D**). A high gastric dysplasia score is equivalent to invasive adenocarcinoma or gastric cancer *in situ* ^8,9,20^. The DKO mice with double deficiency in both TRIF and MyD88 signaling exhibit intermediate dysplasia score. This indicates that deficiency in TRIF signaling mitigates the severe gastric pathology observed in *Myd88*^-/-^ mice in response to *H. felis* infection and therefore, strongly supports our previous studies suggesting that the TRIF pathway plays a role in the development of severe disease pathology. Notably, a recent study citing our previous work suggested that TLR4-TRIF signaling was important in a MALT lymphoma model that involves infection of mice with *H. sius* ^11^. Together, this and our current study indicate that the TRIF signaling pathway and consequently downstream activation of IFN-I pathway may be responsible for the early development and accelerated disease progression thereby supporting a role for downstream target genes of this pathway in promoting disease progression. Indeed, we showed in the current study that upregulation of ISGs was associated with high-grade dysplasia.

**Figure 3.**
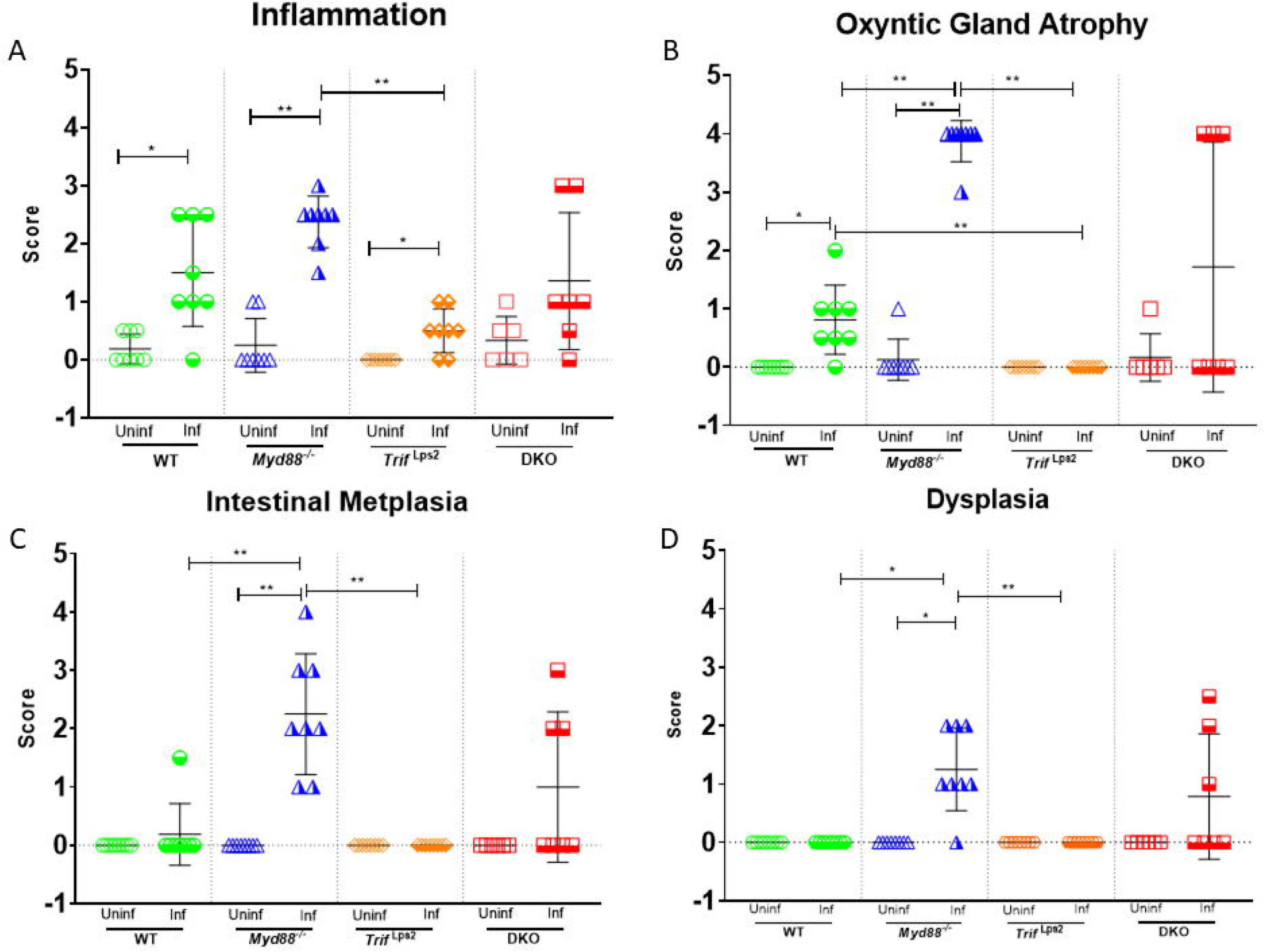
Histopathologic scores in mouse gastric tissue in response to *H. felis* infection. WT and *Myd88*^-/-^ *Trif*^Lps2^, and double knockout (*Myd88^-/-^* and *Trif*^Lps2^, DKO) mice were infected with *H. felis* (Inf.) for 3 months or left uninfected (Uninf.). Mouse stomachs were processed and scored double-blind for histologic disease severity for inflammation (A), gland atrophy (B), intestinal metaplasia (C) and dysplasia (D) on an ascending scale from 0 (no lesions) to 4 (severe lesions).

### Elevated ISG expression in gastric tissue samples from H. felis-infected Myd88^-/-^ mice mirrors ISG expression in patient gastric tissue biopsy samples and is closely associated with the progression of gastric cancer

As ISGs have not previously been implicated in *Helicobacter*-induced disease, we probed gene profiles from previous studies of *H. pylori* induced disease. We reanalyzed microarray data (Gene Expression Omnibus, GEO; Accession number GSE27411)^26^ that contained samples collected from gastric biopsies of *H. pylori*-infected patients with atrophic gastritis. The data set also included gastric biopsies from patients who were negative for *H. pylori* infection ^26^. These data revealed that ISGs that were overexpressed in *Myd88*^-/-^ mice in response to infection with *H. felis* were indeed similarly elevated in gastric tissue samples from *H. pylori*-induced chronic atrophic gastritis patients (**Fig. 4**). Given that chronic atrophic gastritis is deemed as the first step in the multi-step pre-neoplasia progression cascade,^32^ overexpression of ISGs could signal a transition to severe disease pathology. We validated these observations by using available gastric samples obtained from the UCSD’s Biorepository. We found several ISGs significantly upregulated in human gastric cancer biopsy samples that were also upregulated in gastric tissue samples from *H. felis*-infected *Myd88*^-/-^ mice (**Fig. 5**). This finding of similarly highly expressed ISGs between *H. felis*-infected *Myd88*^-/-^ mice and human gastric tissue samples from *H. pylori*-infected and gastric cancer patients validates the mouse data and strongly indicates a role for these ISGs in disease progression to gastric cancer.

**Figure 4.**
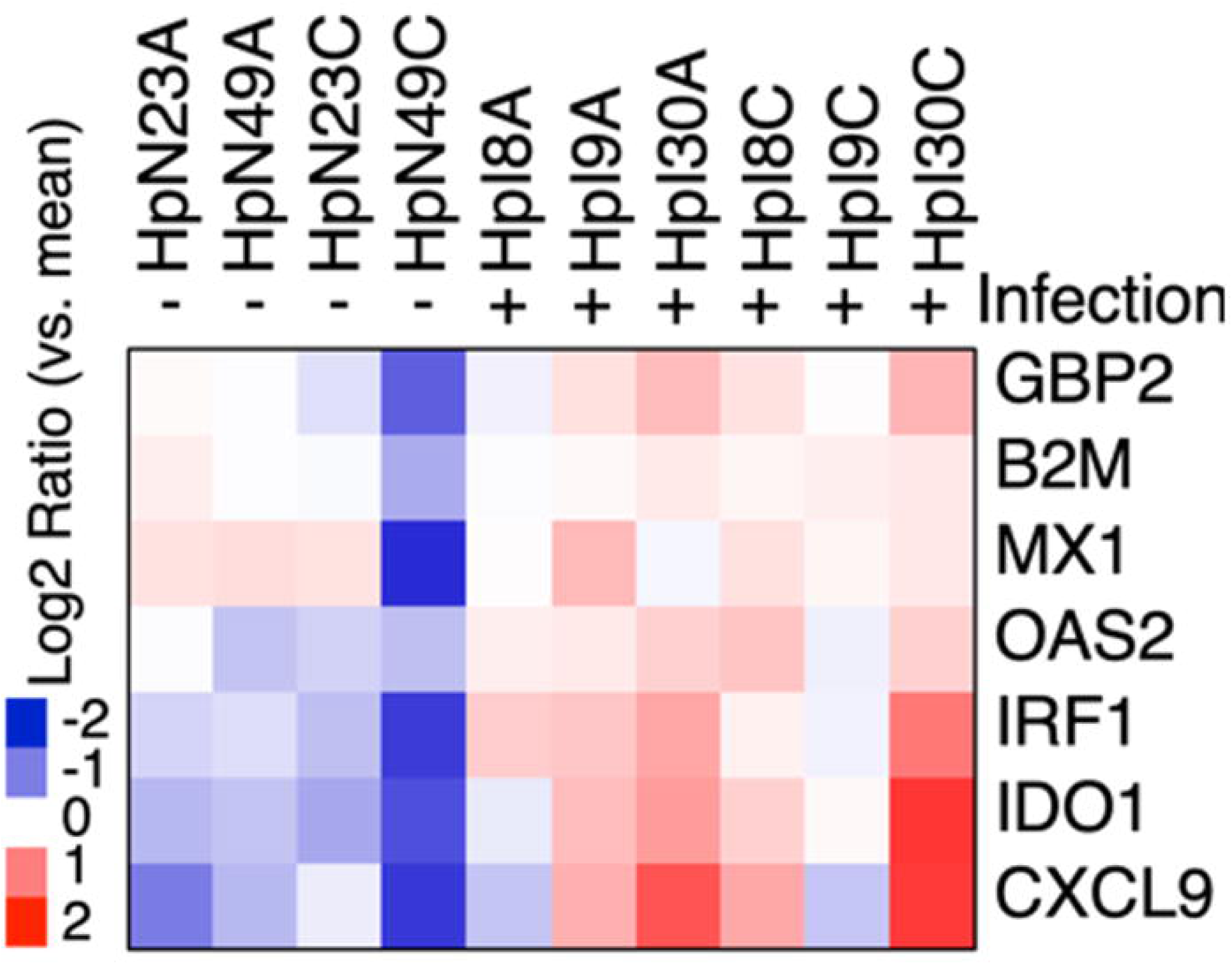
Reanalysis of published microarray data set of gastric tissue from uninfected and *H. pylori*-infected patient with atrophic gastritis^26^ using publicly available dataset (GSE27411).^26^ Samples from *H. pylori*-infected patients are denoted by “+” (n=6) and uninfected by “-“ (n=4).

**Figure 5.**
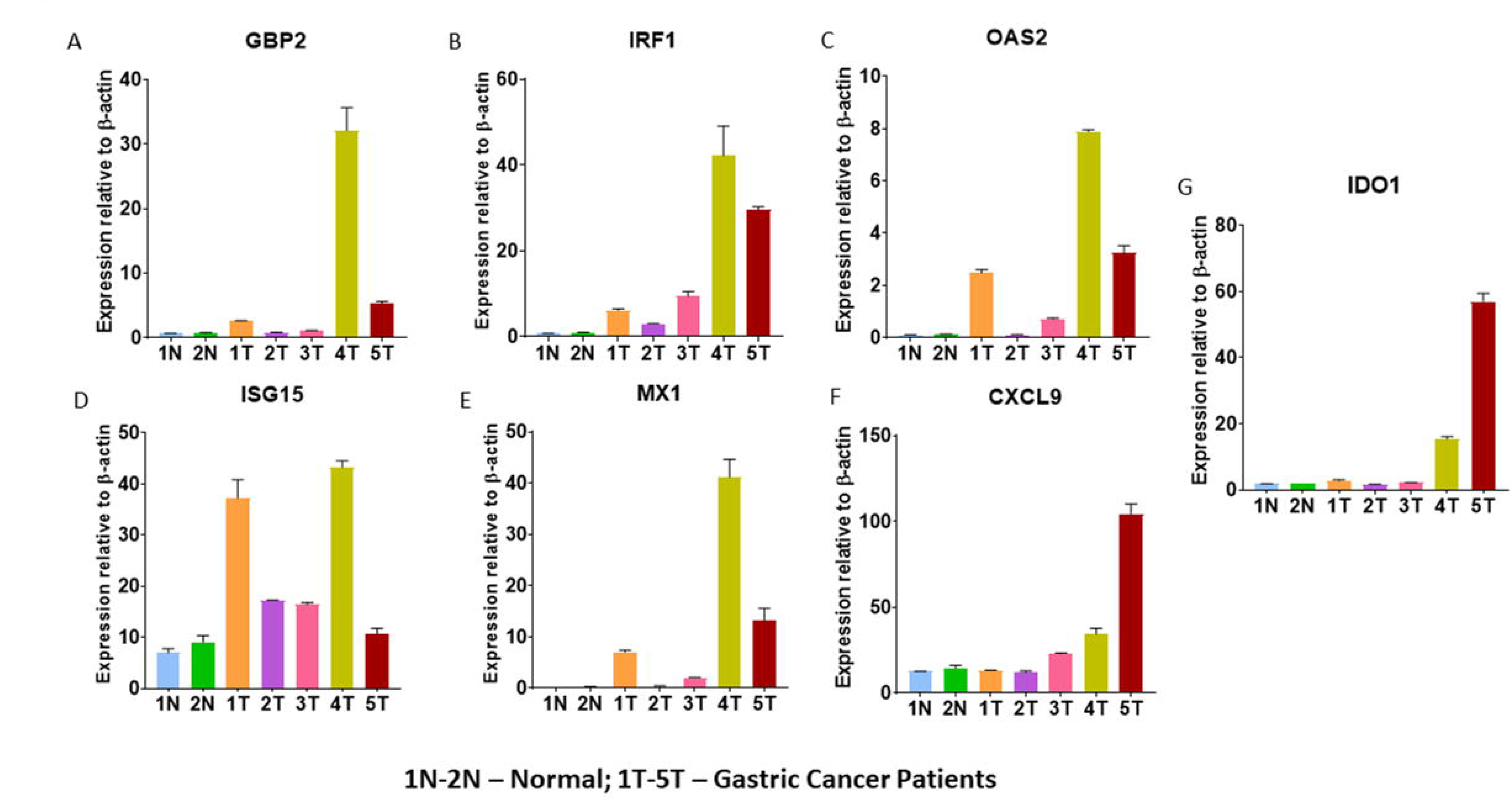
Expression of ISGs, including Gbp2 (A), Irf1 (B), Oas2 (C), Isg15 (D), Mx1 (E), Cxcl9 (F), and Ido1 (G) in gastric cancer patients gastric tissue obtained from the UCSD Biorepository. N, denotes normal and T, tumor.

### Validation of mouse data in gastric cancer patient biopsy samples using Kaplan-Meier survival analysis

In a previous study we found that mice deficient in MyD88 showed rapid progression of gastric pathology upon *H. felis* infection. *Myd88^-/-^* mice developed more severe gastric pathology than WT mice upon infection with *H. felis*, including severe gland atrophy, hyperplasia, intestinal metaplasia, and dysplasia as observed in their stomachs^7^. Two recently published clinical data sets of human stomach cancers from The Cancer Genome Atlas (TCGA)^33,34^ revealed the presence of *Myd88* gene deletions and mutations in gastric and esophageal adenocarcinomas. These findings together with data from our animal model of gastric cancer led us to perform Kaplan-Meier analysis of *Myd88* expression in gastric cancer patients^28^ to validate the clinical relevance of data from our mouse studies. The total number of gastric cancer patients in this database (1,065) ^28^ was divided into high and low expression of *Myd88*. The resulting plots of gastric cancer patients showed that the survival rate of patients with low expression of *Myd88* was significantly lower than that of patients with higher expression in OS (*P* = 0.0016, log-rank test), FP (*P* = 0.0049, log-rank test), and PPS (*P* = 0.022, log-rank test) survival analyses (**Fig. 6**). These results show a link between MyD88 deficiency and gastric cancer and indicate that the level of *Myd88* expression during infection with *Helicobacter* is an important prognosticator of disease outcome.

**Figure 6.**
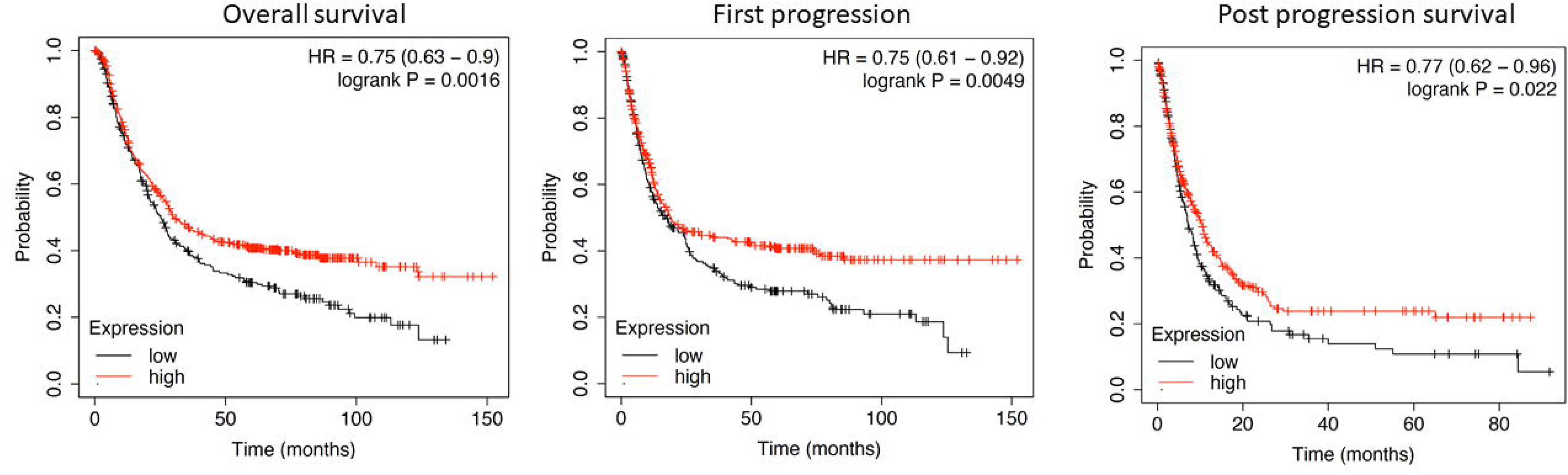
The Kaplan-Meier survival curves for *Myd88* expression in patients with gastric cancer.^28^ This online database included data from 1065 gastric cancer patients. OS, overall survival; FP, first progression; PPS, post-progression survival.

We next analyzed expression of *Trif* in human gastric cancer patients using the Kaplan-Meier survival analysis^28^. Gastric cancer patients with high expression of *Trif* had a significantly lower survival rate than that of patients with lower *Trif* expression in the OS (log-rank *P* = 0.0077), FP (log-rank *P* = 0.0011), and PPS (log-rank *P* = 0.00039) survival analysis. This indicates that high expression of *Trif* is associated with poor prognosis in gastric cancer patients (**Fig. 7**). This supports observations from our mouse studies indicating that hyperactivation of the TRIF pathway and consequently downstream activation of the IFN-I signaling pathway plays a role in the promotion of *Helicobacter*-induced gastric cancer progression. Thus, therefore, intercepting the TRIF signaling pathway could abrogate the development of precancerous lesions.

**Figure 7.**
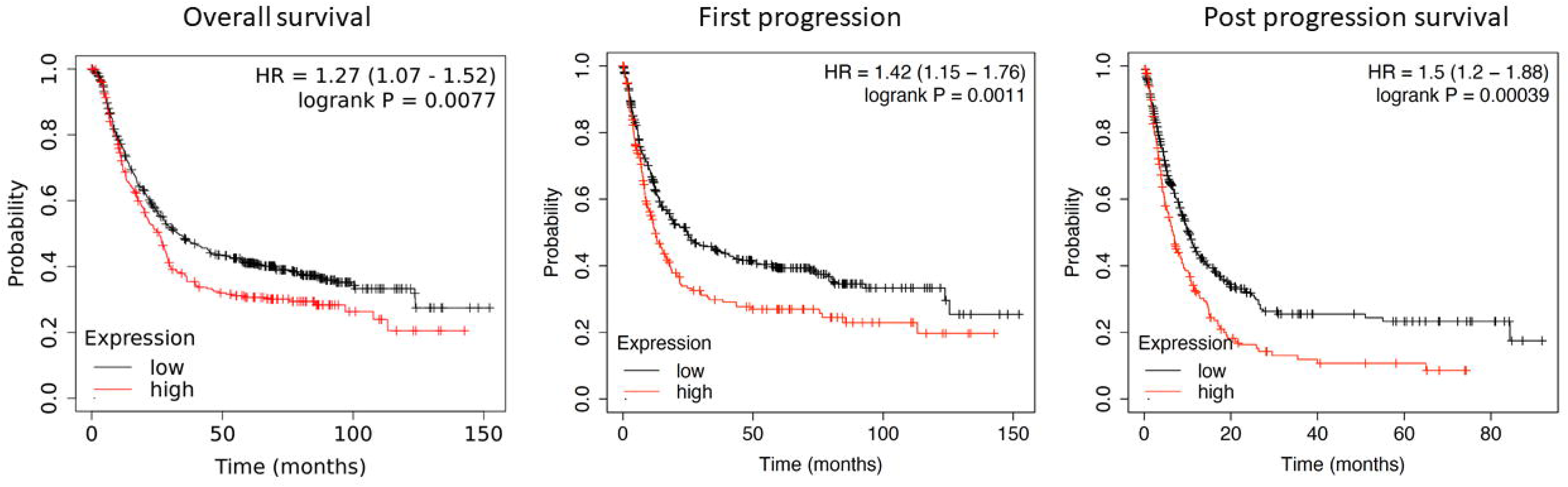
The Kaplan-Meier survival curves for Trif expression in patients with gastric cancer.^28^ This online database included data from 1065 gastric cancer patients. OS, overall survival; FP, first progression; PPS, post-progression survival.

## Discussion

Data from our previous study using *Myd88*^-/-^ mice suggested a link between TRIF activation and *Helicobacter*-induced disease progression to gastric neoplasia^7^. In that study, we showed that mice deficient in MyD88 exhibited rapid progression of gastric pathology following infection with *H. felis*^7^. The mouse stomachs presented severe gland atrophy, hyperplasia, intestinal metaplasia, and dysplasia. However, the molecular mechanism involved in the phenotype remained mostly tenuous. In a separate study, we had previously demonstrated that *H. pylori* induced higher production of IFNα in bone marrow-derived macrophages (BMDM) from *Myd88*^-/-^ mice than those from WT mice,^35^ also strongly indicating that IFN-I response was enhanced in the absence of MyD88. This, therefore, led us to hypothesize that downstream effects of activated IFN-I signaling pathway and target genes (ISGs) played a role in the accelerated disease progression to preneoplastic lesions we observed in *Myd88*^-/-^ mice^7^. Notably, a recent study using a gastric MALT lymphoma model with *H. sius* for infection demonstrated that activation of the TRIF-IFN-I signaling pathway was important for lymphoma development^11^. However, it remained unknown when the gastric epithelial changes associated with this disease progression occurred. In addition, although there was evidence for a role of the TRIF-IFN-I pathway in promoting *Helicobacter*-induced disease progression towards gastric cancer, the role of downstream IFN-I target genes (ISGs)^29^, also remained unknown. Herein, we show that in the absence of MyD88 signaling, changes in the gastric epithelium of mice occurred as early as one month following infection with *H. felis* and that these changes were significant at three months with mice developing preneoplastic lesions. We also show that upregulation of ISGs was associated with severe disease pathology including high-grade dysplasia in mice following infection with *H. felis*, suggesting a potential role for these ISGs in promoting gastric cancer development and progression.

A meaningful observation that emerged from the use of our *H. felis*-infected My88^-/-^ model was that it showed strong parallels to human disease. Specifically, we found that ISGs that were upregulated in this infection model were also upregulated in gastric tissues from *H. pylori*-infected patients with atrophic gastritis (**Fig. 4**) ^26^. Moreover, our analysis of gastric biopsy samples from our UCSD biorepository exhibited upregulation of these ISGs in gastric cancer patients (**Fig. 5**). Together, these data imply that ISGs may be involved in disease transition to malignancy. Upregulation of these ISGs may drive molecular changes associated with *Helicobacter*-induced disease progression to severe gastric pathology, including precancerous lesions.

Not much is known about the role of ISGs in *Helicobacter* induced disease pathogenesis. Some of the ISGs that were upregulated both in mice and patient gastric tissue samples, including *Gbp2* and *Ido1,* have been associated with other human disease conditions. As an example, *Gbp2*^10^ has been shown to be highly expressed in human esophageal squamous cell carcinoma^36^, where its expression is associated with increased proliferation^37^. Notably, this is in line with enhanced proliferation we observed in our *Myd88*^-/-^ infection mouse model ^7^. *Ido1*, another ISG that was highly expressed in gastric tissues of *Myd88*^-/-^ mice following infection with *H. felis*, is suggested to play a role in immune tolerance and high expression is correlated with a poor clinical outcome in multiple cancer types^38–49^. Upregulation of ISG15 expression in colon cancer tissues has been reported to promote proliferation and metastasis and is associated with poor prognosis in colon cancer patients^50^. However, to directly assess the role of specific ISGs in driving this severe disease pathology, targeted inhibition studies are needed to assess more accurately their contribution in disease pathology. Nevertheless, our results strongly indicate that upregulation of ISGs was involved in the severe gastric pathology we observed in *Myd88*^-/-^ mice (**Fig. 1**)^7^ and in human gastric patients (**Fig. 4 & 5**). Cumulatively, our data show that Helicobacter-induced disease progresses rapidly to severe disease pathology in the presence of an unopposed activated TRIF pathway (**Fig. 8**), leading to the upregulation of ISGs.

**Figure 8.**
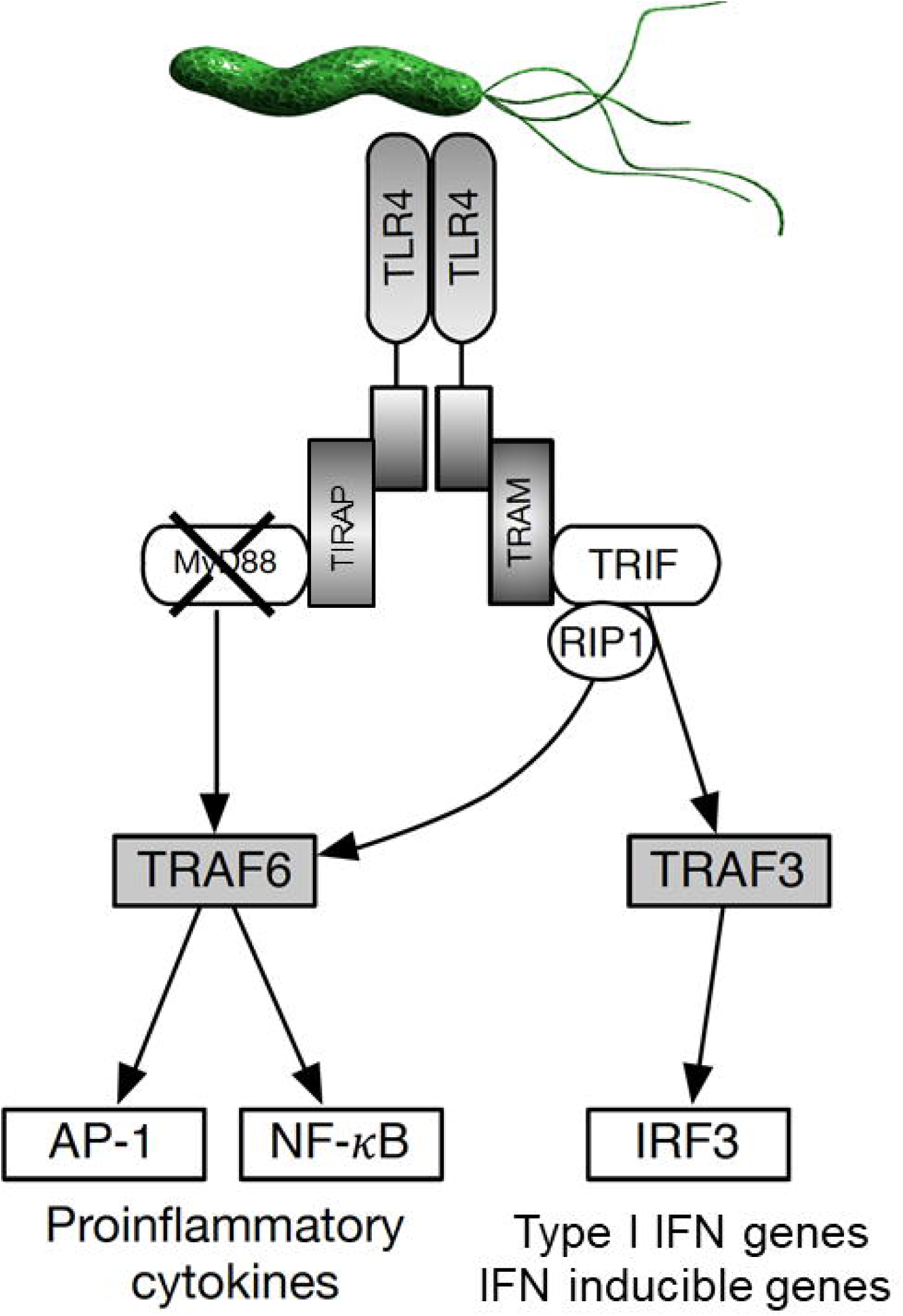
Schematic representation of MyD88 and TRIF signaling pathways during infection of mice with *H. felis* in the absence of MyD88 signaling. In an environment with unopposed activated TRIF pathway disease progresses rapidly leading to upregulation of ISGs, which were associated with severe gastric pathology.

A previous study reported that activation of IFN-I during *Helicobacter* infection contributed to host protection^51^, which contradicts findings from our current study and those of Yamamoto and colleagues^11^, IFN-I signals were mediated via an intracellular protein, nucleotide-binding oligomerization domain 1 (NOD1). In Yamamoto *et al*^11^and our study, activation of IFN-I during infection with *Helicobacter* was mediated by TRIF. The different proteins mediating IFN-I signals could explain the different disease pathogenesis.

In summary, we provide crucial in vivo models and patient-derived evidence that the TRIF-IFN-I pathway is involved in *Helicobacter* disease pathogenesis to gastric cancer. Our findings on the involvement of the TRIF-IFN-I pathway in *Helicobacter*-induced disease progression highlight an important role of this pathway in gastric carcinogenesis, which indicate that this is a crucial area of further exploration. Modulating this pathway in the context of disease progression intervention has the potential to lead to the discovery of novel therapeutic targets. Findings from this study also show that examining mechanisms of *Helicobacter*-induced disease during the early phase of the disease will improve the ability to detect drivers of disease progression given that epithelial changes occur early during the infection. This has the potential to lead to the identification of effective molecular targets that could stop the disease from progressing to the neoplastic lesions and ultimately gastric cancer.

## ACKNOWLEDGMENTS

This work is supported by funding from the Department of Defense (DOD), award W81XWH-20-1-0675 to M.O. The project was also partially funded by NIH grants, UL1TR001442 and P30 DK120515.

## References

1. IARC. Schistomes, liver flukes and Helicobacter pylori. IARC Monographs on the evaluation of carcinogenesis risks to humans. IARC Sci Publ 61, 1–241 (1994).

2. Peek, RM, Jr. & Blaser, MJ. Helicobacter pylori and gastrointestinal tract adenocarcinomas. Nat Rev Cancer 2, 28–37 (2002).

3. SEER. Cancer statistics review (CSR): Stomach cancer.

4. Torre, LA, Siegel, RL, Ward, EM & Jemal, A. Global Cancer Incidence and Mortality Rates and Trends--An Update. Cancer Epidemiol Biomarkers Prev 25, 16–27 (2016).

5. Fox, JG & Wang, TC. Inflammation, atrophy, and gastric cancer. J Clin Invest 117, 60–9 (2007).

6. Correa, P & Piazuelo, MB. The gastric precancerous cascade. J Dig Dis 13, 2–9 (2012).

7. Banerjee, A, Thamphiwatana, S, Carmona, EM, Rickman, B, Doran, KS & Obonyo, M. Deficiency of the myeloid differentiation primary response molecule MyD88 leads to an early and rapid development of Helicobacter-induced gastric malignancy. Infect Immun 82, 356–63 (2014).

8. Lee, CW, Rickman, B, Rogers, AB, Ge, Z, Wang, TC & Fox, JG. Helicobacter pylori eradication prevents progression of gastric cancer in hypergastrinemic INS-GAS mice. Cancer Res 68, 3540–8 (2008).

9. Stenstrom, B, Zhao, CM, Rogers, AB, Nilsson, HO, Sturegard, E, Lundgren, S, Fox, JG, Wang, TC, Wadstrom, TM & Chen, D. Swedish moist snuff accelerates gastric cancer development in Helicobacter pylori-infected wild-type and gastrin transgenic mice. Carcinogenesis 28, 2041–6 (2007).

10. Lozano-Pope, I, Sharma, A, Matthias, M, Doran, KS & Obonyo, M. Effect of myeloid differentiation primary response gene 88 on expression profiles of genes during the development and progression of Helicobacter-induced gastric cancer. BMC Cancer 17, 133 (2017).

11. Yamamoto, K, Kondo, Y, Ohnishi, S, Yoshida, M, Sugiyama, T & Sakamoto, N. The TLR4-TRIF-type 1 IFN-IFN-gamma pathway is crucial for gastric MALT lymphoma formation after Helicobacter suis infection. iScience 24, 103064 (2021).

12. Kawai, T & Akira, S. TLR signaling. Cell Death Differ 13, 816–25 (2006).

13. Hoebe, K, Du, X, Georgel, P, Janssen, E, Tabeta, K, Kim, SO, Goode, J, Lin, P, Mann, N, Mudd, S, Crozat, K, Sovath, S, Han, J & Beutler, B. Identification of Lps2 as a key transducer of MyD88-independent TIR signalling. Nature 424, 743–8 (2003).

14. Oshiumi, H, Matsumoto, M, Funami, K, Akazawa, T & Seya, T. TICAM-1, an adaptor molecule that participates in Toll-like receptor 3-mediated interferon-beta induction. Nat Immunol 4, 161–7 (2003).

15. Yamamoto, M, Sato, S, Hemmi, H, Hoshino, K, Kaisho, T, Sanjo, H, Takeuchi, O, Sugiyama, M, Okabe, M, Takeda, K & Akira, S. Role of adaptor TRIF in the MyD88-independent toll-like receptor signaling pathway. Science 301, 640–3 (2003).

16. Bali, P, Coker, J, Lozano-Pope, I, Zengler, K & Obonyo, M. Microbiome Signatures in a Fast- and Slow-Progressing Gastric Cancer Murine Model and Their Contribution to Gastric Carcinogenesis. Microorganisms 9 (2021).

17. Bali, P, Lozano-Pope, I, Pachow, C & Obonyo, M. Early detection of tumor cells in bone marrow and peripheral blood in a fastprogressing gastric cancer model. Int J Oncol 58, 388–396 (2021).

18. Mejias-Luque, R, Lozano-Pope, I, Wanisch, A, Heikenwalder, M, Gerhard, M & Obonyo, M. Increased LIGHT expression and activation of non-canonical NF-kappaB are observed in gastric lesions of MyD88-deficient mice upon Helicobacter felis infection. Sci Rep 9, 7030 (2019).

19. Obonyo, M, Rickman, B & Guiney, DG. Effects of myeloid differentiation primary response gene 88 (MyD88) activation on Helicobacter infection in vivo and induction of a Th17 response. Helicobacter 16, 398–404 (2011).

20. Rogers, AB, Taylor, NS, Whary, MT, Stefanich, ED, Wang, TC & Fox, JG. Helicobacter pylori but not high salt induces gastric intraepithelial neoplasia in B6129 mice. Cancer Res 65, 10709–15 (2005).

21. Hernandez, J, Turner, MA, Bali, P, Hosseini, M, Bouvet, M, Kelly, K & Obonyo, M. Genomically Silent Refractory Gastric Cancer in a Young Patient Exhibits Overexpression of CXCL5. Current Oncology 29, 4725–4733 (2022).

22. Dobin, A, Davis, CA, Schlesinger, F, Drenkow, J, Zaleski, C, Jha, S, Batut, P, Chaisson, M & Gingeras, TR. STAR: ultrafast universal RNA-seq aligner. Bioinformatics 29, 15–21 (2013).

23. Heinz, S, Benner, C, Spann, N, Bertolino, E, Lin, YC, Laslo, P, Cheng, JX, Murre, C, Singh, H & Glass, CK. Simple combinations of lineage-determining transcription factors prime cis-regulatory elements required for macrophage and B cell identities. Mol Cell 38, 576–89 (2010).

24. Love, MI, Huber, W & Anders, S. Moderated estimation of fold change and dispersion for RNA-seq data with DESeq2. Genome Biol 15, 550 (2014).

25. Zhou, Y, Zhou, B, Pache, L, Chang, M, Khodabakhshi, AH, Tanaseichuk, O, Benner, C & Chanda, SK. Metascape provides a biologist-oriented resource for the analysis of systems-level datasets. Nat Commun 10, 1523 (2019).

26. Nookaew, I, Thorell, K, Worah, K, Wang, S, Hibberd, ML, Sjovall, H, Pettersson, S, Nielsen, J & Lundin, SB. Transcriptome signatures in Helicobacter pylori-infected mucosa identifies acidic mammalian chitinase loss as a corpus atrophy marker. BMC Med Genomics 6, 41 (2013).

27. Saldanha, AJ. Java Treeview--extensible visualization of microarray data. Bioinformatics 20, 3246–8 (2004).

28. Szasz, AM, Lanczky, A, Nagy, A, Forster, S, Hark, K, Green, JE, Boussioutas, A, Busuttil, R, Szabo, A & Gyorffy, B. Cross-validation of survival associated biomarkers in gastric cancer using transcriptomic data of 1,065 patients. Oncotarget 7, 49322–49333 (2016).

29. Schneider, WM, Chevillotte, MD & Rice, CM. Interferon-stimulated genes: a complex web of host defenses. Annu Rev Immunol 32, 513–45 (2014).

30. El-Zimaity, HM, Ota, H, Graham, DY, Akamatsu, T & Katsuyama, T. Patterns of gastric atrophy in intestinal type gastric carcinoma. Cancer 94, 1428–36 (2002).

31. Goldenring, JR & Nam, KT. Oxyntic atrophy, metaplasia, and gastric cancer. Prog Mol Biol Transl Sci 96, 117–31 (2010).

32. Shah, SC, Piazuelo, MB, Kuipers, EJ & Li, D. AGA Clinical Practice Update on the Diagnosis and Management of Atrophic Gastritis: Expert Review. Gastroenterology 161, 1325–1332 e7 (2021).

33. Cancer Genome Atlas Research, N. Comprehensive molecular characterization of gastric adenocarcinoma. Nature 513, 202–9 (2014).

34. Cancer Genome Atlas Research, N, Analysis Working Group: Asan, U, Agency, BCC, Brigham, Women’s, H, Broad, I, Brown, U, Case Western Reserve, U, Dana-Farber Cancer, I, Duke, U, Greater Poland Cancer, C, Harvard Medical, S, Institute for Systems, B, Leuven, KU, Mayo, C, Memorial Sloan Kettering Cancer, C, National Cancer, I, Nationwide Children’s, H, Stanford, U, University of, A, University of, M, University of North, C, University of, P, University of, R, University of Southern, C, University of Texas, MDACC, University of, W, Van Andel Research, I, Vanderbilt, U, Washington, U, Genome Sequencing Center: Broad, I, Washington University in St, L, Genome Characterization Centers, BCCA, Broad, I, Harvard Medical, S, Sidney Kimmel Comprehensive Cancer Center at Johns Hopkins, U, University of North, C, University of Southern California Epigenome, C, University of Texas, MDACC, Van Andel Research, I, Genome Data Analysis Centers: Broad, I, Brown, U, Harvard Medical, S, Institute for Systems, B, Memorial Sloan Kettering Cancer, C, University of California Santa, C, University of Texas, MDACC, Biospecimen Core Resource: International Genomics, C, Research Institute at Nationwide Children’s, H, Tissue Source Sites: Analytic Biologic, S, Asan Medical, C, Asterand, B, Barretos Cancer, H, BioreclamationIvt, Botkin Municipal, C, Chonnam National University Medical, S, Christiana Care Health, S, Cureline, Duke, U, Emory, U, Erasmus, U, Indiana University School of, M, Institute of Oncology of, M, International Genomics, C, Invidumed, Israelitisches Krankenhaus, H, Keimyung University School of, M, Memorial Sloan Kettering Cancer, C, National Cancer Center, G, Ontario Tumour, B, Peter MacCallum Cancer, C, Pusan National University Medical, S, Ribeirao Preto Medical, S, St. Joseph’s, H, Medical, C, St. Petersburg Academic, U, Tayside Tissue, B, University of, D, University of Kansas Medical, C, University of, M, University of North Carolina at Chapel, H, University of Pittsburgh School of, M, University of Texas, MDACC, Disease Working Group: Duke, U, Memorial Sloan Kettering Cancer, C, National Cancer, I, University of Texas, MDACC, Yonsei University College of, M, Data Coordination Center, CI & Project Team: National Institutes of, H. Integrated genomic characterization of oesophageal carcinoma. Nature 541, 169–175 (2017).

35. Obonyo, M, Sabet, M, Cole, SP, Ebmeyer, J, Uematsu, S, Akira, S & Guiney, DG. Deficiencies of myeloid differentiation factor 88, Toll-like receptor 2 (TLR2), or TLR4 produce specific defects in macrophage cytokine secretion induced by Helicobacter pylori. Infect Immun 75, 2408–14 (2007).

36. Guimaraes, DP, Oliveira, IM, de Moraes, E, Paiva, GR, Souza, DM, Barnas, C, Olmedo, DB, Pinto, CE, Faria, PA, De Moura Gallo, CV, Small, IA, Ferreira, CG & Hainaut, P. Interferon-inducible guanylate binding protein (GBP)-2: a novel p53-regulated tumor marker in esophageal squamous cell carcinomas. Int J Cancer 124, 272–9 (2009).

37. Gorbacheva, VY, Lindner, D, Sen, GC & Vestal, DJ. The interferon (IFN)-induced GTPase, mGBP-2. Role in IFN-gamma-induced murine fibroblast proliferation. J Biol Chem 277, 6080–7 (2002).

38. Corm, S, Berthon, C, Imbenotte, M, Biggio, V, Lhermitte, M, Dupont, C, Briche, I & Quesnel, B. Indoleamine 2,3-dioxygenase activity of acute myeloid leukemia cells can be measured from patients’ sera by HPLC and is inducible by IFN-gamma. Leuk Res 33, 490–4 (2009).

39. Creelan, BC, Antonia, S, Bepler, G, Garrett, TJ, Simon, GR & Soliman, HH. Indoleamine 2,3-dioxygenase activity and clinical outcome following induction chemotherapy and concurrent chemoradiation in Stage III non-small cell lung cancer. Oncoimmunology 2, e23428 (2013).

40. Ferns, DM, Kema, IP, Buist, MR, Nijman, HW, Kenter, GG & Jordanova, ES. Indoleamine-2,3-dioxygenase (IDO) metabolic activity is detrimental for cervical cancer patient survival. Oncoimmunology 4, e981457 (2015).

41. Inaba, T, Ino, K, Kajiyama, H, Yamamoto, E, Shibata, K, Nawa, A, Nagasaka, T, Akimoto, H, Takikawa, O & Kikkawa, F. Role of the immunosuppressive enzyme indoleamine 2,3-dioxygenase in the progression of ovarian carcinoma. Gynecol Oncol 115, 185–92 (2009).

42. Ino, K, Yamamoto, E, Shibata, K, Kajiyama, H, Yoshida, N, Terauchi, M, Nawa, A, Nagasaka, T, Takikawa, O & Kikkawa, F. Inverse correlation between tumoral indoleamine 2,3-dioxygenase expression and tumor-infiltrating lymphocytes in endometrial cancer: its association with disease progression and survival. Clin Cancer Res 14, 2310–7 (2008).

43. Speeckaert, R, Vermaelen, K, van Geel, N, Autier, P, Lambert, J, Haspeslagh, M, van Gele, M, Thielemans, K, Neyns, B, Roche, N, Verbeke, N, Deron, P, Speeckaert, M & Brochez, L. Indoleamine 2,3-dioxygenase, a new prognostic marker in sentinel lymph nodes of melanoma patients. Eur J Cancer 48, 2004–11 (2012).

44. Wainwright, DA, Balyasnikova, IV, Chang, AL, Ahmed, AU, Moon, KS, Auffinger, B, Tobias, AL, Han, Y & Lesniak, MS. IDO expression in brain tumors increases the recruitment of regulatory T cells and negatively impacts survival. Clin Cancer Res 18, 6110–21 (2012).

45. Ino, K, Yoshida, N, Kajiyama, H, Shibata, K, Yamamoto, E, Kidokoro, K, Takahashi, N, Terauchi, M, Nawa, A, Nomura, S, Nagasaka, T, Takikawa, O & Kikkawa, F. Indoleamine 2,3-dioxygenase is a novel prognostic indicator for endometrial cancer. Br J Cancer 95, 1555–61 (2006).

46. Okamoto, A, Nikaido, T, Ochiai, K, Takakura, S, Saito, M, Aoki, Y, Ishii, N, Yanaihara, N, Yamada, K, Takikawa, O, Kawaguchi, R, Isonishi, S, Tanaka, T & Urashima, M. Indoleamine 2,3-dioxygenase serves as a marker of poor prognosis in gene expression profiles of serous ovarian cancer cells. Clin Cancer Res 11, 6030–9 (2005).

47. Ricciuti, B, Leonardi, GC, Puccetti, P, Fallarino, F, Bianconi, V, Sahebkar, A, Baglivo, S, Chiari, R & Pirro, M. Targeting indoleamine-2,3-dioxygenase in cancer: Scientific rationale and clinical evidence. Pharmacol Ther (2018).

48. Brochez, L, Chevolet, I & Kruse, V. The rationale of indoleamine 2,3-dioxygenase inhibition for cancer therapy. Eur J Cancer 76, 167–182 (2017).

49. Ferdinande, L, Decaestecker, C, Verset, L, Mathieu, A, Moles Lopez, X, Negulescu, AM, Van Maerken, T, Salmon, I, Cuvelier, CA & Demetter, P. Clinicopathological significance of indoleamine 2,3-dioxygenase 1 expression in colorectal cancer. Br J Cancer 106, 141–7 (2012).

50. Zhou, S, Ren, M, Xu, H, Xia, H, Tang, Q & Liu, M. Inhibition of ISG15 Enhances the Anti-Cancer Effect of Trametinib in Colon Cancer Cells. Onco Targets Ther 12, 10239–10250 (2019).

51. Watanabe, T, Asano, N, Fichtner-Feigl, S, Gorelick, PL, Tsuji, Y, Matsumoto, Y, Chiba, T, Fuss, IJ, Kitani, A & Strober, W. NOD1 contributes to mouse host defense against Helicobacter pylori via induction of type I IFN and activation of the ISGF3 signaling pathway. J Clin Invest 120, 1645–62 (2010).

