## Supplementary Material for "Activation of the TRIF pathway and downstream targets results in the development of precancerous lesions during infection with *Helicobacter*"

Table of contents	page numbers

Table S1	2

Table S2	3

Supplementary Figure legends	4

Fig. S1	5

**Table S1. Characteristics of gastric cancer patients.**

| **Patient ID** | **Patient Sex** | **Patient Age** | **Patient race/ethnicity** | **Primary** | **Grade** | **Metastatic** | **Stage** | **Chemotherapy** |
| --- | --- | --- | --- | --- | --- | --- | --- | --- |
| 1T | F | 25 | Hispanic | Adenocarcinoma, diffuse type | G3: poorly diff | Yes | IV (ypT4bypN3bypM1) | EOX/FOLFIRI |
| 2T | M | 66 | Asian | Adenocarcinoma, residual | G3: poorly diff | No | IIA (ypT3N0) | EOX and chemorads with capecitabine |
| 3T | M | 51 | White | Adenocarcinoma | G3: poorly diff | No | IIB (ypT4aN0) | Yes (unspecified in notes) |
| 4T | M | 78 | White | Invasive adenocarcinoma | G3: poorly diff | Yes | IIIC (pT4aN3a) | No |
| 5T | M | 69 | Hispanic | adenocarcinoma | G3: poorly diff | Yes | IV (ypT4bN3bM1) | FOLFOX |

T; tumor gastric cancer tissue, EOX; epirubicin, oxaliplatin, and capecitabine, FOLFOX; folic acid, fluorouracil, and oxaliplatin

**Table S2: Primers Used for RT-PCR analysis:**

| **Primer name** | **Sequence (5´-3´)** | **Reference** |
| --- | --- | --- |
| hOAS2-F \| GCTTCCGACAATCAACAGCCAAG; hOAS2-R \| CTTGACGATTTTGTGCCGCTCG | | This study |
| hIDO1-F \| GCCTGATCTCATAGAGTCTGGC; hIDO1-R \| TGCATCCCAGAACTAGACGTGC | | This study |
| hISG15-F \| TCCTGCTGGTGGTGGACAA; hISG15-R \| TTGTTATTCCTCACCAGGATGCT | | ^1^ |
| hMX1-F \| GTGCATTGCAGAAGGTCAGA; hMX1-R \| TCAGGAGCCAGCTTAGGTGT | | ^1^ |
| hGBP2-F \| GATTTCACCCTGGAACTGGA; hGBP2-R \| GGGTTCAGCTCTTCCTCCTT | | ^2^ |
| hβactin-F \| GATTACTGCTCTGGCTCCTAGC; hβactin-R \| GACTCATCGTACTCCTGCTTGC | | ^2^ |
| hIRF1-F \| CTGTGCGAGTGTACCGGATG; hIRF1-R \| ATCCCCACATGACTTCCTCTT | | ^3^ |
| hCXCL9-F \| CCAGTAGTGAGAAAGGGTCGC; hCXCL9-R \| AGGGCTTGGGGCAAATTGTT | | ^3^ |

1	Liu, B. C. *et al.* Constitutive Interferon Maintains GBP Expression Required for Release of Bacterial Components Upstream of Pyroptosis and Anti-DNA Responses. *Cell Rep* **24**, 155-168 e155 (2018). <https://doi.org:10.1016/j.celrep.2018.06.012>

2	Qin, A. *et al.* Guanylate-binding protein 1 (GBP1) contributes to the immunity of human mesenchymal stromal cells against Toxoplasma gondii. *Proc Natl Acad Sci U S A* **114**, 1365-1370 (2017). <https://doi.org:10.1073/pnas.1619665114>

3	Spandidos, A., Wang, X., Wang, H. & Seed, B. PrimerBank: a resource of human and mouse PCR primer pairs for gene expression detection and quantification. *Nucleic Acids Res* **38**, D792-799 (2010). <https://doi.org:10.1093/nar/gkp1005>

**Supplementary Figure legends**

Histopathologic scores of uninfected mouse tissues sections from WT, *Myd88*^-/-^, *Trif^Lps2^*, and DKO mice at 1, 3, or, 6 months. Mouse stomachs tissue sections were scored for histologic disease severity on an ascending scale from 0 (no lesions) to 4 (severe lesions) for inflammation (A), epithelial defects (B), Oxyntic gland atrophy (C), Hyperplasia (D), Intestinal metaplasia (E), Dysplasia (F).
